# Alpha-synuclein null mutation exacerbates the phenotype of a model of Menkes disease in female mice

**DOI:** 10.1101/2023.11.15.567255

**Authors:** MegAnne Casey, Dan Zou, Renee A. Reijo Pera, Deborah E. Cabin

## Abstract

Genetic modifier screens provide a useful tool, in diverse organisms from *Drosophila* to *C. elegans* and mice, for recovering new genes of interest that may reduce or enhance a phenotype of interest. This study reports a modifier screen, based on N-ethyl-N-nitrosourea (ENU) mutagenesis and outcrossing, designed to increase understanding of the normal function of murine α-synuclein (*Snca*). Human *SNCA* was the first gene linked to familial Parkinson’s disease. Since the discovery of the genetic link of *SNCA* to Parkinson’s nearly three decades ago, numerous studies have investigated the normal function of SNCA protein with divergent roles associated with different cellular compartments. Understanding of the normal function of murine Snca is complicated by the fact that mice with homozygous null mutations live a normal lifespan and have only subtle synaptic deficits. Here, we report that the first genetic modifier (a sensitized mutation) that was identified in our screen was the X-linked gene, *ATPase copper transporting alpha (Atp7a).* In humans, mutations in *Atp7a* are linked to to Menkes disease, a disease with pleiotropic phenotypes that include a severe neurological component. *Atp7a* encodes a trans-Golgi copper transporter that supplies the copper co-factor to enzymes that pass through the ER-Golgi network. Male mice that carry a mutation in *Atp7a* die within 3 weeks of age regardless of *Snca* genotype. In contrast, here we show that *Snca* disruption modifies the phenotype of *Atp7a* in female mice. Female mice that carry the *Atp7a* mutation, on an *Snca* null background, die earlier (prior to 35 days) at a significantly higher rate than those that carry the *Atp7a* mutation on a wildtype *Snca* background ATPase copper transporting alpha. Thus, *Snca* null mutations sensitize female mice to mutations in *Atp7a,* suggesting that Snca protein may have a protective effect in females, perhaps in neurons, given the co-expression patterns. Although data has suggested diverse functions for human and mouse α-synuclein proteins in multiple cell compartments, this is the first demonstration via use of genetic screening to demonstrate that Snca protein may function in the ER-Golgi system in the mammalian brain in a sex-dependent manner.

**Author summary:** This study sought to probe the normal function(s) of a protein associated with Parkinson’s disease, the second most common neurodegenerative disease in humans. We used a genetic modifier approach to uncover aspects of normal protein function, via mutagenesis of mice and screening for neurological problems that are decreased or enhanced in mice that are null for α-synuclein (*Snca)*. Through these studies, we identified the X-linked gene that is mutated in Menkes disease in humans as a modifier of the null *Snca* phenotype, specifically in female mice. The gene mutated in Menkes disease, *ATP7a*, encodes a copper transporter that is known to act in the trans-Golgi sub-cellular compartment. Genetic modifier effects suggest that Snca may also play a role in that compartment, potentially in the mammalian brain.

## Introduction

Parkinson’s disease (PD) is the second most common neurodegenerative disease in humans with Alzheimer’s disease being the most common^12^. While the majority of cases are late-onset and sporadic, genetic forms of PD have also been identified.^13-15^ Alpha-synuclein (*SNCA*) was the first gene identified as causing a genetic form of the disease^16, 17^, and SNCA protein was subsequently shown to be a major component of Lewy bodies^19^, intracellular inclusions commonly observed in postmortem midbrain tissue in conjunction with PD. The basis of SNCA toxicity in PD is not well understood; human mutations that have been identified include both loss-of-function (rare missense mutations) and gain-of-function (numerical variant) mutations^20-23^. One potential mechanism for toxicity includes a slow build-up of a distinct wildtype or mutant oligomer;^24^ however, the mechanism and identity of the toxic form is not known and continues to be a subject of debate^23, 25-28^. Clearly, SNCA can form a variety of oligomeric structures, including the fibrils found in Lewy bodies. These inclusions themselves, however, may be toxic or protective.

Numerous studies have reported on diverse functions of SNCA in different cellular compartments including the synapse, mitochondria, nucleus, endoplasmic reticulum, and cytoplasm. Functional analysis is complicated by the fact that mice that lack SNCA protein are overall healthy and live a normal lifespan with only subtle synaptic phenotypes^29-31^. Over-expression of wild type human SNCA has been shown to rescue the phenotypes of mice lacking the synaptic chaperone cysteine string protein suggesting a direct or indirect interaction between these proteins^32^. In other studies, Snca has also been shown to enhance SNARE assembly at the synapse in mice and additional mechanisms of toxicity, including inhibition of ER to Golgi trafficking, have also been proposed^23, 25-28, 33-36^. Other functions of SNCA, in mitochondrial dynamics and structure, regulation of gene expression, epigenetic programming, nuclear transport, neuron survival, cytoskeletal stabilization and DNA repair, have also been reported (**Table 1**).

**Table 1.**
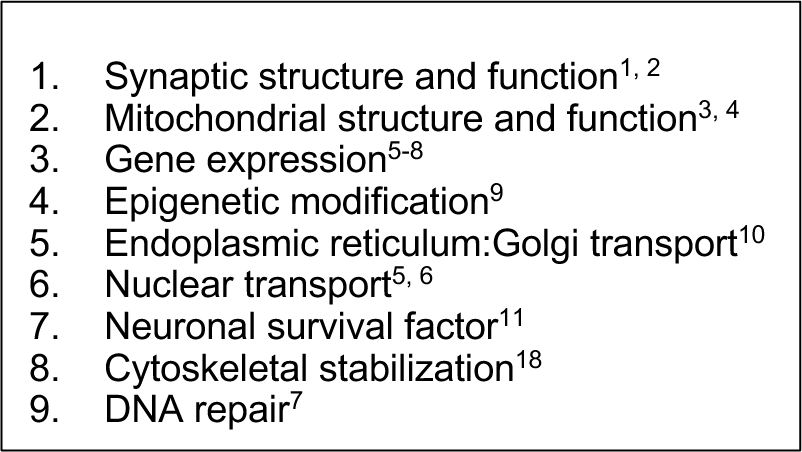
Reported α-Synuclein Functions.

To Here, we probed the function of SNCA further, we sought to use a unique approach relative to other studies and focus on modifiers of the phenotypes association with *Snca* null alleles in mice. We designed a sensitized mutagenesis screen in mice as previously reviewed^37, 38^. The approach of using modifier screening for sensitizing mutation(s) has the advantage of being less biased regarding *a priori* assumptions such as Snca function, cellular and subcellular localization, and physical properties, relative to other common methods of identifying interactions^38, 39^. to the goal of these studies was to identify mice with ENU-induced mutations that result in neurological phenotypes that are more severe in the absence of Snca than in the presence of Snca protein. These ENU-induced mutations would provide provide information on genetic pathways in which Snca functions.

## Results

Our ENU (N-ethyl-N-nitrosourea) screen followed a standard protocol: C57BL/6 male mice were mutagenized with ENU, then bred to females homozygous null for *Snca,* and offspring were outcrossed in a breeding scheme designed to uncover recessive mutations that are more severe in the absence of Snca than on a wild type background (“sensitized”; see **Figure 1**). Approximately 30-40 G3 offspring per line were assessed for neurological phenotypes with 125 pedigrees screened through the G3 stage. Our screen for neurological phenotypes was based on the modified SHIRPA (SmithKline Beecham, Harwell, Imperial College, Rogyal London Hospital, phenotype assessment) protocol established in a large ENU-mutagenesis screen over two decades ago^40^. Seven of the 125 pedigree lines were not eliminated as analyses revealed: 1) Extreme reaction to clicker with clasping of forepaws, 2) poor performance on bar, 3) circling behavior, 4) over reaction to clicker, hyperactivity and odd gait, 5) premature dropping from bar, 6) hard landing, bouncing and little reaction in clicker testing, and 7) poor performance on bar with clasping of hindpaws.

**Figure 1.**
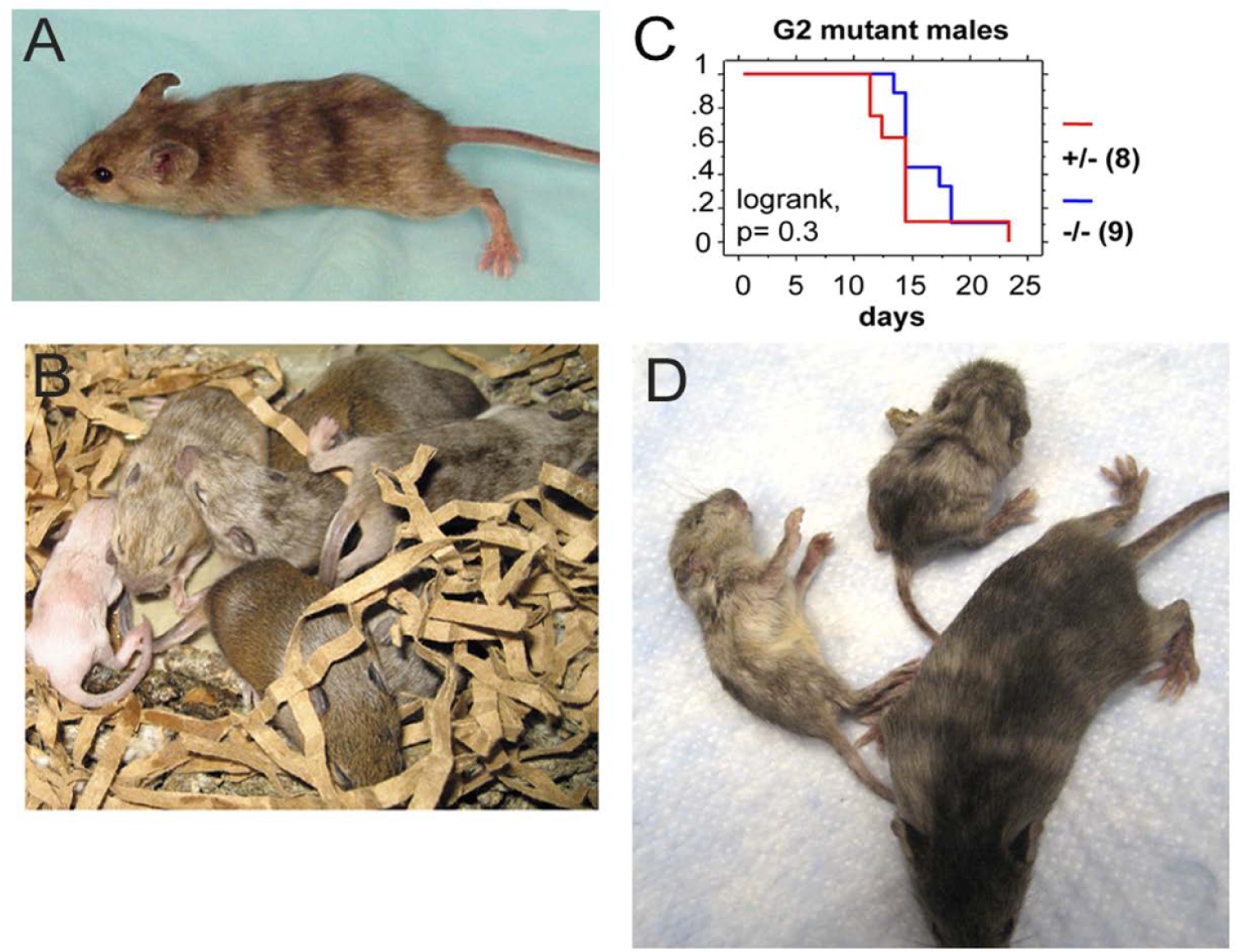
The mouse coat color mutation is X-linked. **A)** One of 2 original G1 females identified as carrying a coat color mutation. **B)** Litter of G2 pups showing a white male, two females with mottled coat color and agouti littermates. **C)** Kaplan-Meyer lifespan analysis shows that lack of Snca does not affect lifespan of coat color mutant males. **D)** *Snca* null, coat color mutant female G3 littermates at 4 weeks of age, showing one viable and two dying animals.

In the second pedigree of the screen, two pheno-deviant G1 females were identified by their patchy coat color (**Figure 1A**). The patchiness suggested X-linkage of the underlying mutation. We surmised that a good candidate, X-linked gene was *Atp7a*, a trans-Golgi copper transporter implicated in neurodegeneration. Indeed, mutations in *ATP7a* are known to cause X-linked Menkes disease in humans, a disease with a severe neurological component. The G1 female phenodeviants were heterozygous for the *Snca* null allele and thus, were bred to *Snca* null mice to determine if the mutation was indeed X-linked, and whether lack of *Snca* affected the severity of the phenotype. A litter is shown in **Figure 1B**; the coat color of mutant males was all white while affected females had a patchy coat color, further confirming X-linkage. We observed that all coat color mutant males died <25 days; further analysis of life span also indicated that there was no difference in lifespan between *Snca* null and *Snca* heterozygotes of the mutant males (**Figure 1C**). In contrast to observations with male coat color mutant mice, female coat color mutants that were also homozygous *Snca* null mutants demonstrated increased early death (<35 days) than did mice heterozygous or wildtype for *Snca*, though statistical significance was not reached with the low number of affected female offspring from the two G1 females (data not shown). Subsequently, G2 coat color mutant females were bred to both *Snca* nulls and to wildtype controls (on the 129S6 genetic background) to obtain greater numbers for comparisons. The G3 offspring demonstrated a significantly higher rate of early death (<35 days) in the *Snca* null coat color females vs *Snca* heterozygotes or wildtype (**Table 2**). Thus, we confirmed that the coat color mutation is sensitized by the null mutation in *Snca*. An example of viable and inviable Snca null coat color mutant female littermates is shown (**Figure 1D).**

**Table 2.** G3 coat color mutant females dead by 35 days.

| **Table 2.** G3 coat color mutant females dead by 35 days |  |  |
| --- | --- | --- |
| **Snca** | **Dead** | **Viable** |
| +/-, +/+ | 9 | 38 |
| -/- | 23 | 20 |
| p=0.00069, X<sup>2</sup> test |  |  |
| **Snca** | **Dead** | **Viable** |
| +/- | 5 | 22 |
| +/+ | 4 | 16 |
| p=1, Fisher's exact test |  |  |

As *Atp7a* was the best candidate gene, a mutant female’s entire *Atp7a* coding region DNA was sequenced. One mutation was found at nucleotide position1951 (NM_001109757) in exon 7 (**Figure 2A, B**), which was heterozygous for G and T. The G changes the amino acid at position 610 from an isoleucine to a serine (NP_001103227; this corresponds to human amino acid 618, NP_000043). More than 400 nonsense, missense and insertions/deletions have been idenitifed in the human *ATP7a* gene; those associated with Menkes disease are within the protein coding sequence and map to the copper associated domains, ATPase or transmembrane domains in particular^41^. In the two parental strains used in the mutagenesis protocol, 129S6 and C57BL6/J, the nucleotide at position 1951 is a T (**Figure 2C, D**). The change to a G at this position segregated with the coat color phenotype; sequences from an affected male and a second affected female are shown (**Figure 2E, F)**. Amino acid 610 lies between the 6^th^ of six metal binding domains in *Atp7a* and the first transmembrane domain, a position of functional importance based on previous studies^42, 43^.

**Figure 2.**
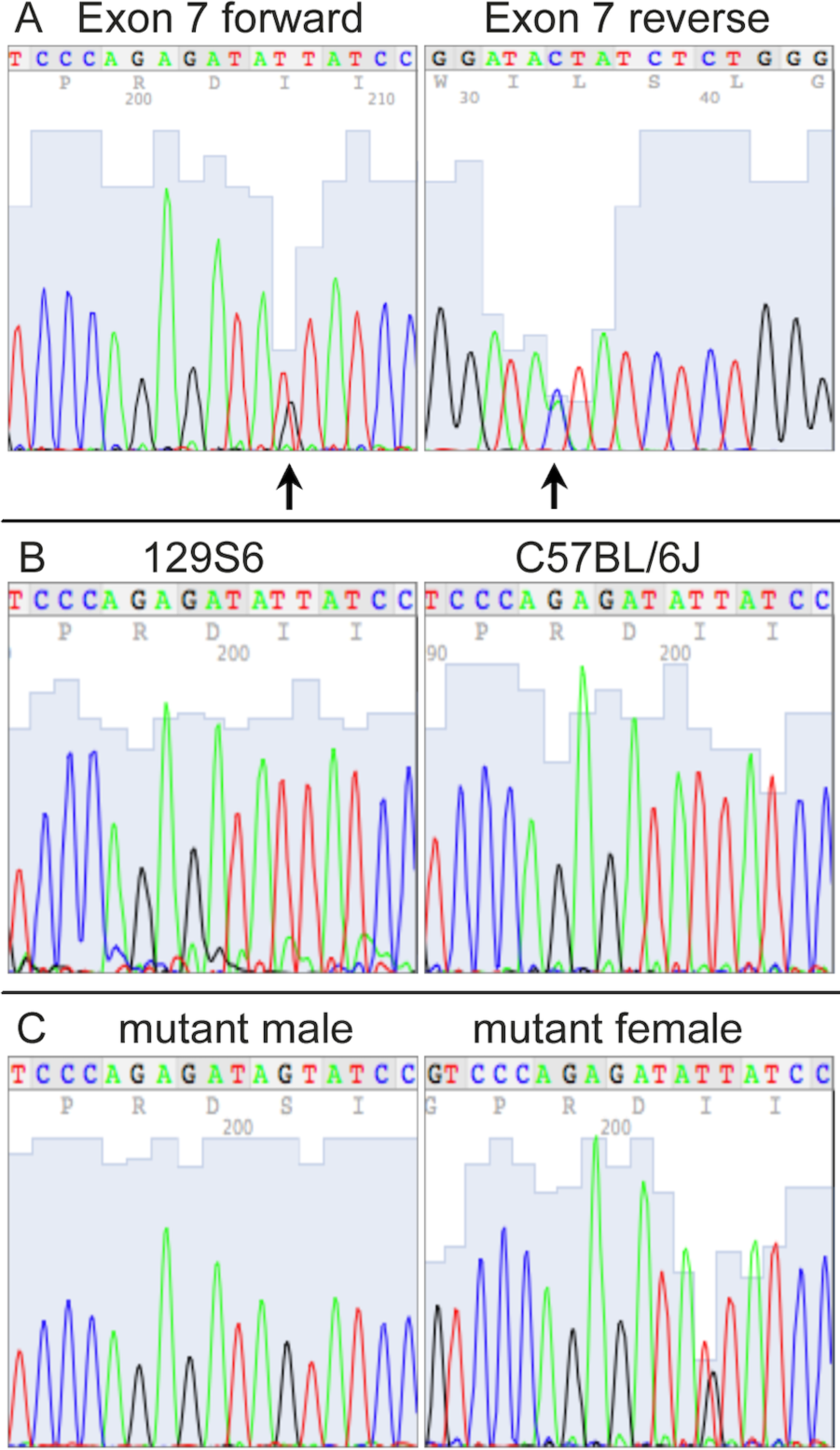
Identification of a mutation in *Atp7a* that segregates with the coat color phenotype. **A)** Forward and reverse exon 7 sequence from a coat color mutant female identifies coding sequence nucleotide 1951 as heterozygous (arrows). **B)** Exon 7 forward sequence from females of the two strains used in the ENU mutagenesis procedure, 129S6 and C57BL/6J. **C)** Exon 7 forward sequence from an affected male is hemizygous for the mutant nucleotide, and an additional affected female is heterozygous.

A multi-species protein alignment of exon 7 is shown (**Figure 3A)**. The isoleucine at position 610 is well conserved, though a conservative substitution of a valine is found in both the opossum and zebrafish. Numerous *Atp7a* alleles are known in mice, and most cause affected males to die *in utero*^44^. The observation that males carrying the I610S mutation do not die until two to three weeks postnatally suggests that this new allele may be hypomorphic resulting in only partial loss-of-function. Northern blot analysis indicated similar amounts of *Atp7a* mRNA in both male and female mutant brains (data not shown). Similarly, Western blot analysis with total brain lysates from postnatal day 8 pups showed that Atp7a protein is made in all mutant animals, though at a lower level than in wildtype animals. Further, the absence of Snca protein did not alter Atp7a levels (**Figure 3B**). Similar results were observed in postnatal day 14 animals (data not shown).

**Figure 3.**
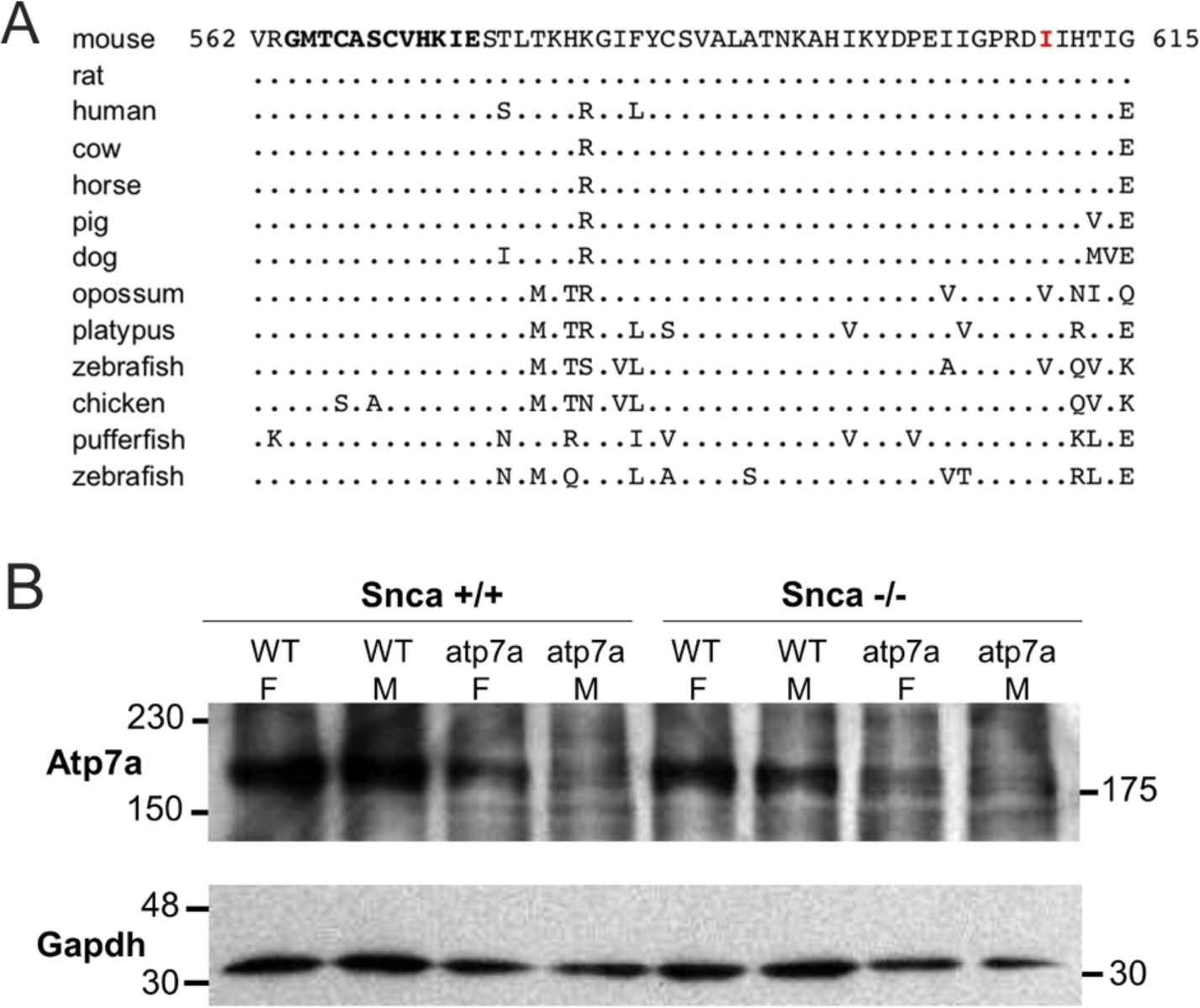
Isoleucine 610 is highly conserved, but Atp7a protein is still produced in the I610S mutant mice. **A)** Alignment of exon 7 amino acid sequences. The mutated I610 is indicated in red, and the sixth Atp7a copper-binding domain is in bold towards the N-terminus. The consensus copper-binding domain is GMT/HCxxCxxxIE. **B)** Western blot of total brain lysates from P8 mice, with Gapdh as a loading control.

Snca is most abundant in neurons and thus we reasoned, it may have a neural protective effect in females. Aneurysms caused by faulty collagen maturation were a frequent cause of death in *Atp7a* male mutant mice that survive postnatally, and Snca clearly did not appear to impact death or survival. In females, random X-inactivation of the mutant chromosome could circumvent phenotypic problems associated with collagen dysfunctionat least in part. We hypothesized that while random X-inactivation would also occur in neurons, some neuronal populations might be more vulnerable to the effects of mutant Atp7a in the absence of Snca. Thus, to test our hypothesis, we compared brain samples from viable and dying *Snca*-/-; *Atp7a^I610S^* female littermates via an antibody against cleaved caspase-3 to identify apoptotic cells (**Figure 4**). The cerebral cortex appeared to be smaller relative to the cerebellum in brains from dying mice (**Figure 4A, inset**), and patches of cortical neurons positive for cleaved caspase-3 were observed only in dying but not in viable female littermates with the *Snca*-/-;*Atp7a^I610S^*genotype (**Figure 4A, B**). In addition to cleaved caspase-3 positivity as an indicator of apoptosis, haematoxylin counterstain also demonstrated patches of large numbers of smaller pyknotic nuclei in cortical layers II/III in dying but not in viable females (**Figure 4B, C**).

**Figure 4.**
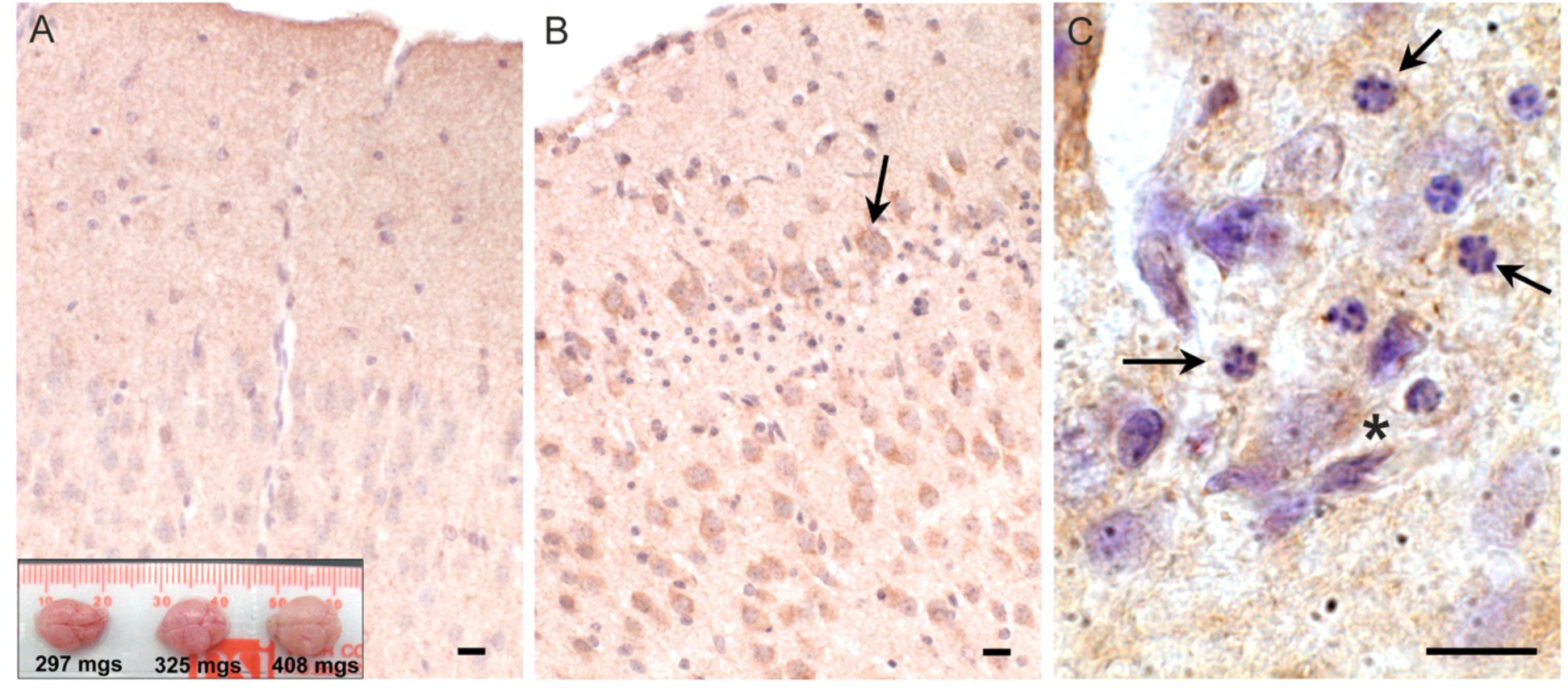
Apoptosis in cerebral cortex of inviable *Atp7a^I610S^* Snca null female mice. Cleaved caspase-3 immunohistochemistry. **A)** Cingulate cortex from viable *Snca-/-; Atp7a^I610S^* female shown in Fig. 1A; inset; brains from the three animals in Fig. 1A, viable on the right. **B)** Cingulate cortex from inviable *Snca-/-; Atp7a^I610S^*female; arrow indicates cleaved-caspase-3 positive neuron above layer of pyknotic nuclei. **C)** Higher magnification of pyknotic nuclei (arrows) and cleaved caspase-3 positive neuron (asterisk). Scale bars = 0.05mm.

Atp7a immunohistochemistry was performed on brains from male mice in which only one allele of *Atp7a* was expressed. Sections of postnatal day fifteen brains from *Atp7a^I610S^*males were reduced in size relative to age-matched controls, so sections of postnatal day eight control brains of similar size were also analyzed by immunohistochemistry. Greater numbers of pyknotic nuclei were detected in cerebral cortex, mostly in layers II/III, in *Atp7a* mutant brains relative to controls (**Figure 5A-D**). Immunohistochemical staining of Atp7a was strong in the corpus callosum in fifteen-day old control animals (**Figure 5E**). Staining was weaker in 15 day mutant males (**Figure 5F, G**), but the 15 day *Snca*+/+;*Atp7a^I610S^* pattern more closely resembled that of the 8 day control (**Figure 5H**) than does the 15 day *Snca*-/-;*Atp7a^I610S^*male. Analysis of *Snca*-/-;*Atp7a^I610S^*brain samples suggested a delay in axonal localization of Atp7a, while the *Snca*+/+;*Atp7a^I610S^*brain samples had more diffuse staining similar to the 8 day controls. Atp7a staining of cell bodies is more apparent in the fifteen-day old mutants and the eight day controls than in the fifteen day control brain.

**Figure 5.**
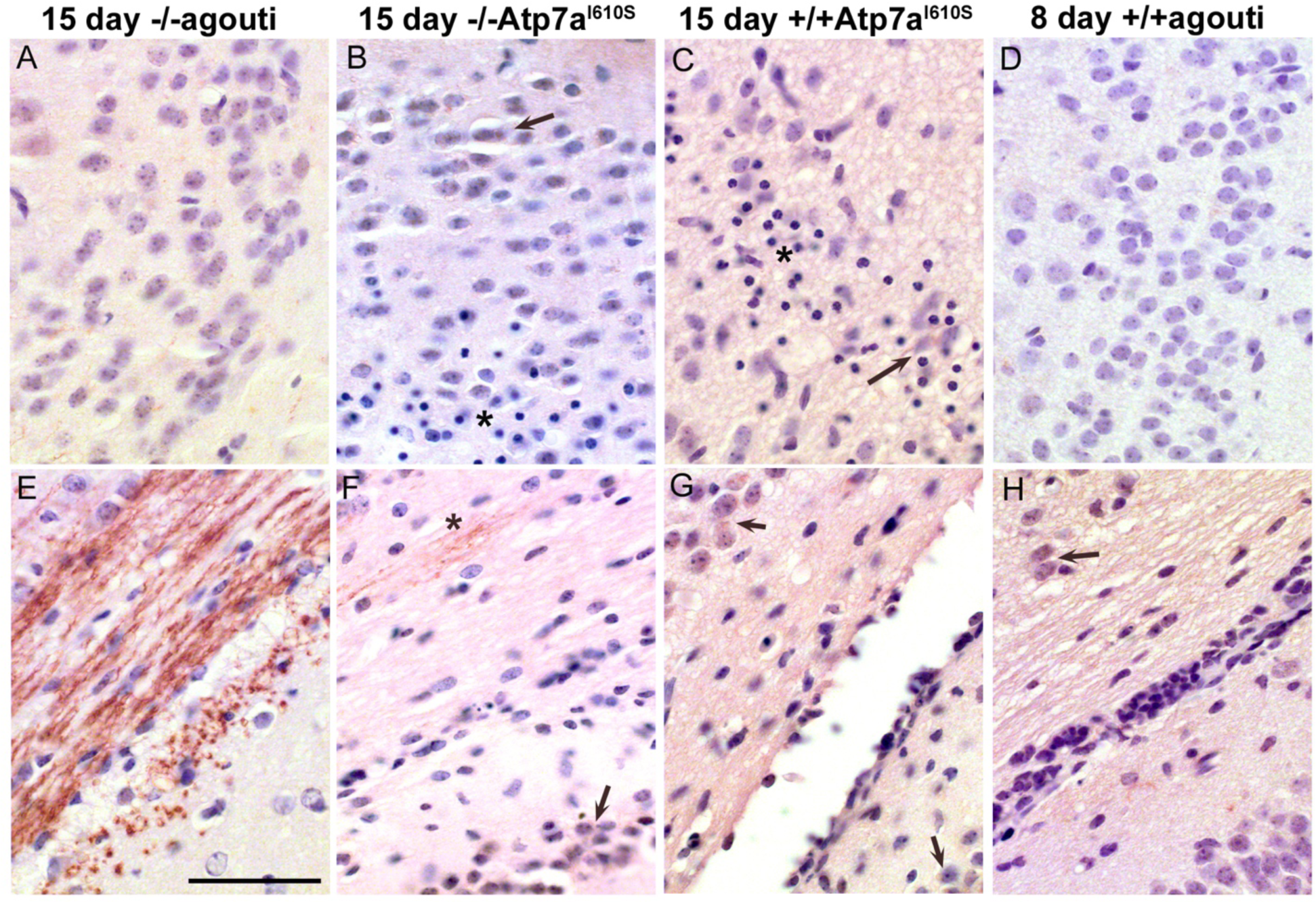
*Atp7a* in male control and mutant cerebral cortex and corpus callosum. *Atp7a* immunohistochemistry on brains of *Snca-/-; Atp7a^I610S^*, *Snca+/+; Atp7a^I610S^*, and control animals at 15 days of age, as well as an 8 day control to match brain size of the 15 day old mutants. **A-D)** Cerebral cortex. In mutant animals, **B** and **C**, arrows indicate cell body localization of *Atp7a*; asterisks indicate the regions with pyknotic nuclei. **E-H)** Corpus callosum. **E)** At 15 days, *Atp7a* is strongly expressed in wild type. **F)** Asterisk indicates minor neuritic-like *Atp7a* localiztion. **F, G,** and **H**) Arrows indicate cell body localization of *Atp7a*. Scale bar for all panels = 0.05mm.

Strong *Atp7a* staining was observed in the developing striatum in the fifteen-day control brain (**Figure 6A**), in a neuritic pattern. **Figure 6A** shows an *Snca* null brain with staining similar to *Snca* wild type brain samples (data not shown). The defined pattern of Atp7a staining in axonal sub-compartments of the striatum was not established in the corresponding region from fifteen day mutant and eight day old control brains (**Figure 6C-H**). In the *Snca*-/-;*Atp7a^I610S^*brain, Atp7a was localized primarily to cell bodies (**Figure 6 D**). However, in the presence of Snca, some faint neuritic staining was apparent in fifteen-day mutant brain samples (**Figure 5E, F**), although cell body staining was more prominent than in the fifteen-day control. In the absence of Snca (on an *Snca* null background), faint staining of wild type Atp7a was detected in a neuritic pattern in eight-day old brain. Thus, Snca may aid in the proper localization of the mutant form of Atp7a, but is not required by the wild type protein.

**Figure 6.**
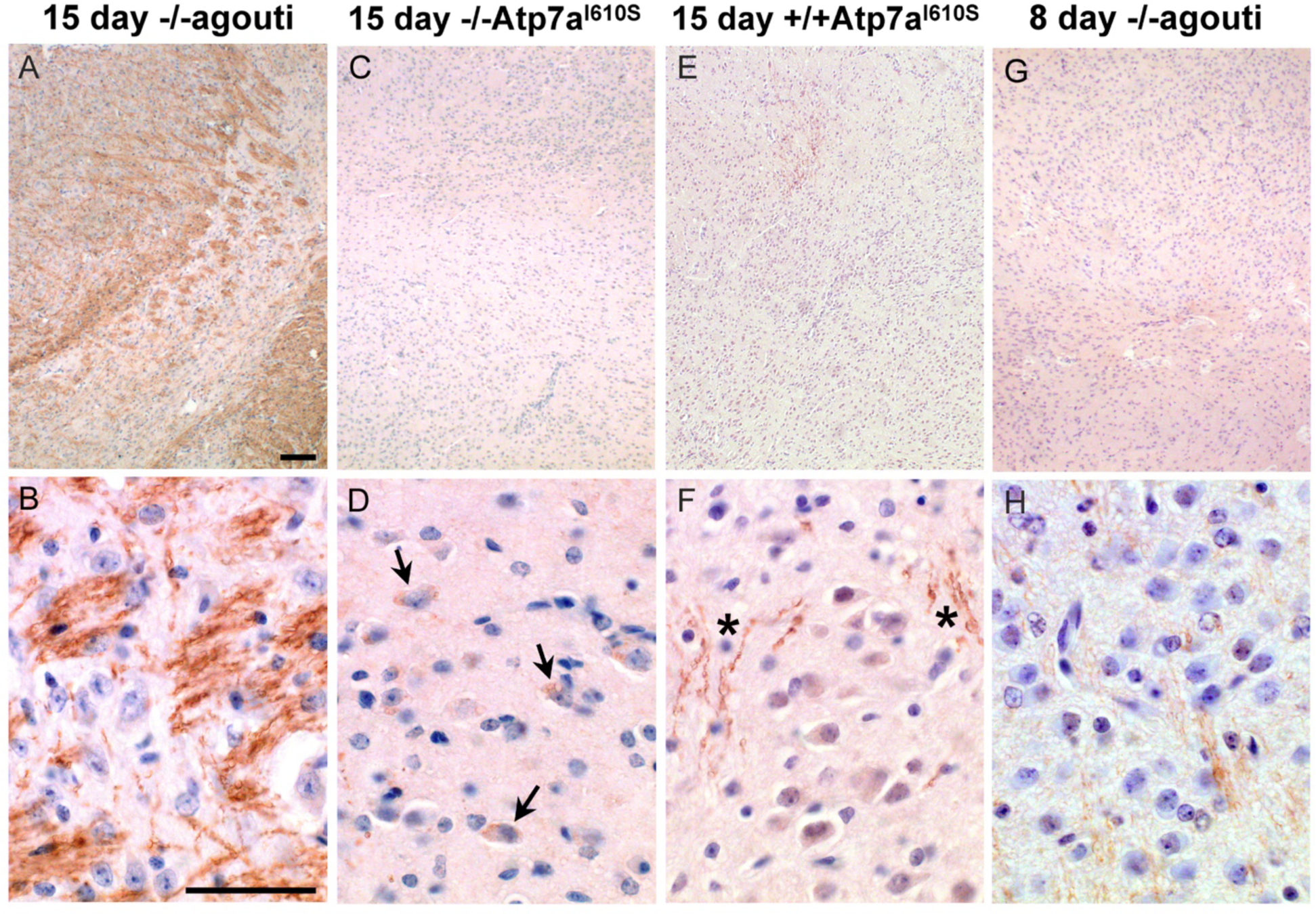
*Atp7a* in developing striatum. *Atp7a* immunohistochemistry on striatum from male mice. Top panels, low magnification, scale bar = 0.1mm; bottom panels high magnification, scale bar = 0.05 mm. **A, B)** Wild type control shows robust Atp7a expression in striatum. **C-F)** 15 day mutant males lag in development of the striatum and appear similar to the 8 day control in panels **G** and **H**. Arrows in **D** indicate cell body localization of Atp7a in the *Snca-/-; Atp7a^I610S^* animal; asterisks in **F** show some neuritic staining in the *Snca+/+; Atp7a^I610S^*animal.

Neuritic localization of Atp7a was unexpected. However, Atp7a has been reported to play a role in axonal targeting and synaptogenesis in olfactory bulb^45-47^. We used immunohistochemistry with trans-Golgi marker, Tgoln2, in double labeling experiments to determine if neuritic Atp7a was associated with the trans-Golgi compartment. In wildtype striatum from 22-day old mice, Atp7a and Tgoln2 showed a similar pattern of localization (**Figure 7A-C**). Atp7a signal was almost completely co-localized with the Tgoln2 signal (**Figure 7A**). Similar results were seen in axonal tracts of the wildtype corpus callosum connecting the hemispheres (not shown). In *Atp7a^I610S^*brains, the Atp7a signal was faint in corpus callosum, but overlapped with that of Tgoln2 (not shown). SH-SY5Y cells were then tested to confirm that the antibodies used were indeed targeting proteins of the proper compartment. In these undifferentiated cells, the antibodies against ATP7a and TGOLN2 both identify a perinuclear region that is consistent with the Golgi compartment (**Figure 6D-F**). Higher magnification confocal imaging indicated that the two proteins are found in the same perinuclear compartment (**Figure 6G-I**).

**Figure 7.**
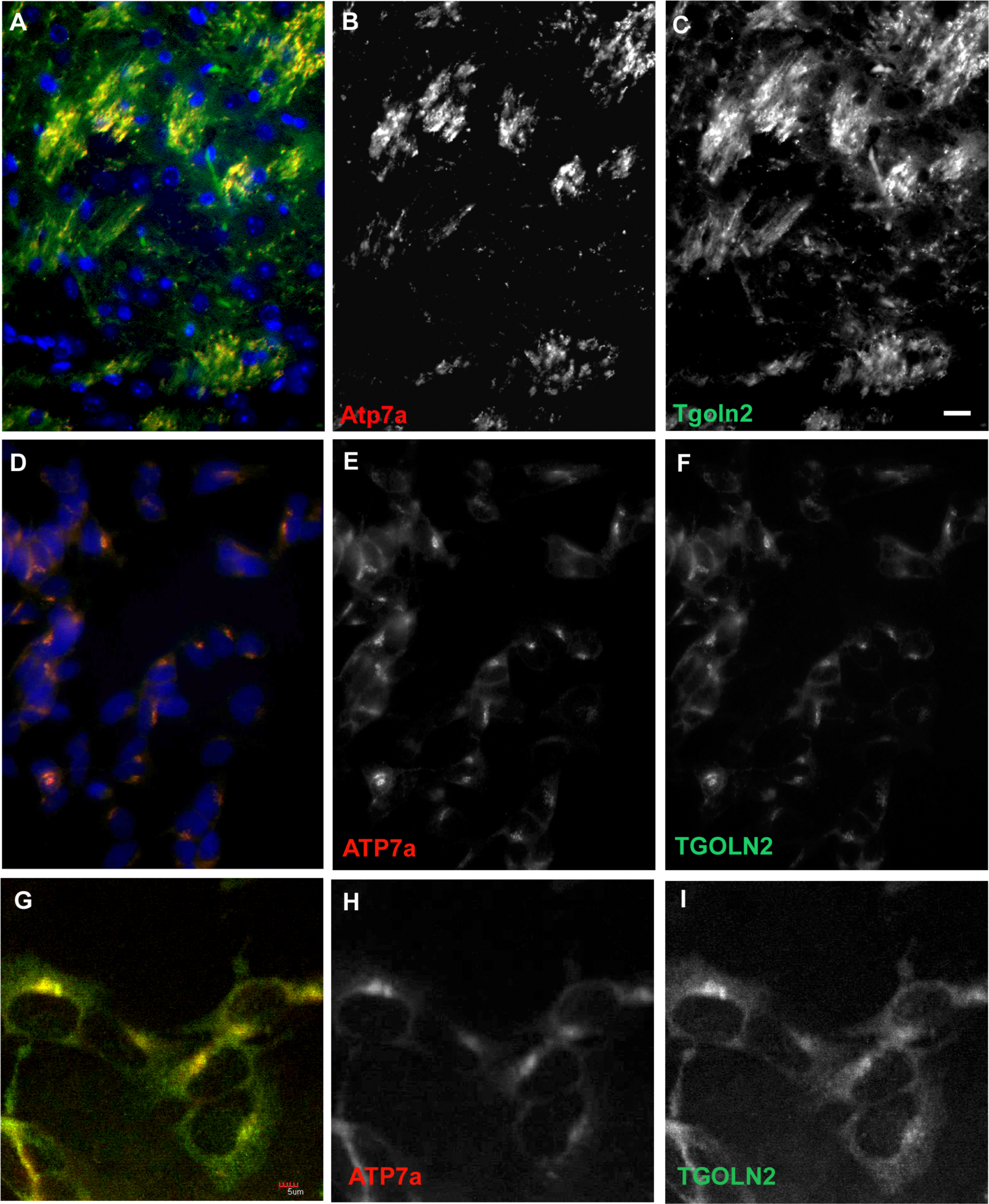
A trans-Golgi marker protein, *Tgoln2*, co-localizes with *Atp7a* in mouse brain and in SH-SY5Y cells. **A-C**) Striatum in 22 day old wild type mouse. **A)** Merge, with DAPI in blue. **B)** *Atp7a*, C, *Tgoln2*. **D–F**) SH-SY5Y cells express *ATP7a* and *TGOLN2* in the same peri-nuclear compartment. **D**) Merge, with DAPI in blue. **E**) *ATP7a*. **F)** *TGOLN2*. Scale bar for **A–F** = 0.05 mm. **G – I**) Single slice confocal image of SH-SY5Y cells. **G**) Merge. **H)** *ATP7a*. **I**) *TGOLN2*. Scale bar for **G–I** = 0.005 mm.

## Discussion

Prior efforts to understand the normal function of Snca have mostly focused on the synapse, though Snca can be found in other cellular compartments such as the nucleus ^48^. We have found that a mutation in *Atp7a* produces a more severe phenotype in the absence of Snca. Atp7a is a well-characterized trans-Golgi copper transporter, thus genetically linking Snca to a function in that compartment in mammalian brain. Previous work has shown that SNCA retards ER to Golgi transport^33, 35^. However, in non-vertebrates in which there are no synucleins, the possibility exists that in the absence of a synaptic compartment, SNCA mislocalizes and causes a phenotype that would never be seen in mammalian neurons. More recent work involving PC12 cells transfected with mutant *SNCA* has shown that the protein has a deleterious effect on SNARE assembly^35^. Again, these cells are not neurons and normally express little SNCA. This is in contrast to our findings of a potential protective effect of Snca on a mutated trans-Golgi protein, an effect that may involve aiding proper localization of the mutated protein. This finding highlights an advantage of the unbiased genetic approach of sensitized mutagenesis screens.

The first pheno-deviant mice identified in our screen were two female littermates identified by a patchy coat color phenotype. The patchy coat color suggested X-linkage, and the best X-linked candidate gene was *Atp7a*, a trans-Golgi copper transporter mutated in Menkes disease in humans (OMIM 309400). This disease affects many organ systems, including the nervous system (see review^49^). All enzymes that pass through the ER-Golgi pathway and require copper as a co-factor are impaired. These include tyrosinase, causing pigmentation defects, and lysyl oxidase, which is required for collagen maturation. Two neuronal enzymes that require copper are dopamine β-hydroxylase, which converts dopamine to norepinephrine, and peptidyglycine α-amidating monooxygenase, required for neuropeptide amidation, a post-translational modification that many neuropeptides require for full activity^50^.

We decided to test whether lack of Snca had any effect on the severity of the phenotypes in our mouse coat color mutants for several reasons. First, Menkes disease has a severe neurological component that can progress to complete decerebration. Second, SNCA has been shown to affect ER-Golgi trafficking in yeast and mammalian cells^35, 51^, and third, occasional reports that SNCA may bind copper^52, 53^. In addition, it has been shown that the incidence of PD is higher amongst melanoma patients, linking SNCA to pigmentation (see review^54^). We found that lack of Snca significantly increased incidence of early death in female coat color mutant mice carrying a missense mutation in *Atp7a.* These findings are the first demonstration in mammalian brain of a functional link between Snca and the trans-Golgi compartment and also link sex-specific effects to *Snca* function.

Potential mechanisms of protective effects of Snca should be examined in the context of the *Atp^I610S^* mutation. We suggest that lack of Snca may impact localization of *Atp7a^I610S^* by affecting its transport from the ER to the Golgi compartment, thus preventing its further transport to the trans-Golgi via Possible mechanisms include two trans-Golgi mechanisms: inefficient SNARE assembly impeding vesicular transport between the two compartmentsor Snca activity as a chaperone to stabilize mutant Atp7a and facilitate trafficking to the trans-Golgi. Further studies of primary cortical neurons from Atp7a mutant male mice may distinguish between these possibilities.

Menkes disease has a strong neurological component, as does Parkinson’s disease. The loss of neuronal sub-types is not well understood. However, this work, and other recent studies, suggest that there are shared etiologies and molecular mechanisms underlying different neurodegenerative disorders. Further examination of the interactions of *Atp7a* and *Snca* in Parkinson’s disease in humans and mice are of interest. Examination of their localization and activity levels in animal models of Parkinson’s disease might reveal an ER-Golgi transport deficit in that disease.

## Methods

### ENU mutagenesis

C57BL/6J males were treated with ENU^37^, allowed to recover fertility, and mated to *Snca* null females on a 129S6 background^30^. G1 males were used in further breeding to uncover sensitized recessive mutations. The *Atp7a* mutation described here arose in 2 G1 females of the same pedigree. Further backcrosses to produce G2 and G3 generations were to 129S6 males that were Snca null or wild type. *Snca* genotyping primers have been described^30^. All procedures on mice have been approved by the McLaughlin Research Institute IACUC, and the Institute’s Animal Resource Center is AAALAC accredited. Statistical analyses were performed with the StatView statistics package (SAS Institute, Cary, NC).

### Sequencing

Primers for sequencing the coding sequence of *Atp7a* (23 amplicons, 22 coding sequence exons) were chosen using the UCSC Genome browser (http://genome.ucsc.edu) link from mouse *Atp7a* to ExonPrimer (http://ihg.helmholtz-muenchen.ed/cgi-bin/primer/ExonPrimerUCSC), with default settings. Sequencing was performed at the UC Davis Sequencing facility (http://www.davissequencing.com/). Exon 7 primers used for assessing the I610S mutation are F:TAAGGCAATCCTGTGCTACG and R: TGATTCCAGAAGGTGGTTGAC, with an amplicon size of 162 bp.

### Western blots

Western blots were performed using whole brain lysates as described^30^. The primary antibodies used were chicken anti-*Atp7a* (Sigma), mouse anti-*Gapdh* (Millipore), and rabbit anti-neuron-specific enolase (Polysciences). Secondary hydrogen peroxidase-conjugated antibodies, goat-anti-chicken, goat-anti-mouse, and goat-anti-rabbit were obtained from GE HealthCare/Amersham, as was ECL Plus for chemiluminescent detection.

### Immunohistochemistry and immunofluorescence

Brains perfused with 4% formaldehye were dehydrated, embedded in paraffin, and cut in 10um sections. Immunohistochemistry was performed as described (Cabin et al., 2005), using the primary antibodies rabbit anti-cleaved caspase-3 (Trevigen) and chicken anti-*Atp7a* (Sigma), Meyer’s haematoxylin was obtained from Sigma. Biotinylated secondary antibodies and VectaStain were from Vector Laboratories. Immunofluorescence was performed as described^55^. Primary antibodies were chicken anti-*Atp7a* as above and rabbit anti-*TGOLN2* (AbCam). Secondary antibodies were Alexa-488 anti-rabbit (Invitrogen) and DyLight-549 anti-chicken (Jackson ImmunoResearch). SH-SY5Y cells were obtained from ATTC.

### Microscopy

Bright-field and immunofluorescence images were captured using a Zeiss Axio-Imager microscope; confocal microscopy was performed on an Olympus FV-1000.

## Acknowledgements

We thank Chris Ebeling for performing the ENU injections, and Dr. George Carlson for developing the sensitized ENU mutagenesis protocol used at McLaughlin Research Institute, and for suggestions to the manuscript. We thank the staff of the Institute’s Animal Resource Center for expert animal care and for their valuable expertise with mouse phenotyping. Renovations to and equipment in the Institute’s Animal Resource Center were funded by major grants and donations from Montana’s Department of Commerce, the M.J. Murdock Charitable Trust, the Browning-Kimball Foundation, the Oakland Family, Sletten Construction, Ian and Nancy Davidson, and others. This work was supported in part by NIH NINDS - R01NS062121 (DEC).

